# Observation of female-male mounting in the carrion crow

**DOI:** 10.1101/2020.05.26.116004

**Authors:** Claudia A.F. Wascher, Friederike Hillemann

## Abstract

In the biological sciences, sexual behaviours in non-human animals are traditionally investigated in the context of reproduction and direct fitness benefits. While the evolutionary functions of non-conceptive sexual behaviours (‘socio-sexual’) remain less well explored, these interactions and displays have been suggested to be important for shaping and maintaining social relationships. Here, we report an observation of a captive female carrion crow, *Corvus corone corone*, mounting her co-housed male partner. We highlight the importance of more systematic research and discussions of rarely observed behaviours in social evolution research, including considerations for behaviours that transcend binary or heteronormative frameworks, to provide a more comprehensive understanding of non-conceptive socio-sexual behaviours.

## Introduction

Non-conceptive socio-sexual behaviours, such as same-sex mountings, female-male mountings, or ritualised greetings involving sexual gestures are primarily considered to contribute to managing social relationships and have been described in a variety of species across taxa (Bailey and Zuk, 2009; Gómez et al., 2023; MacFarlane et al., 2010). While also discussed as ‘behavioural errors’ due to mistaken identity (*e.g*., Macchiano et al., 2018), several hypotheses have been proposed regarding these behaviours’ adaptive social functions (Ham et al., 2023 and references therein), including sexual solicitation (Gunst et al., 2020), practice (Manson et al., 1997), play (Lilley et al., 2020), dominance assertion (Faraut et al., 2015), appeasement in socially tense situations (Manson et al., 1997), social bonding (Dal Pesco and Fischer, 2018) and alliance formation (Kotrschal et al., 2006). A well-described behaviour in mammals are social mounts (same-sex mounts or female-male mounts; Dagg, 1984), which may be part of ritualised greetings and may involve genital manipulation (*e.g*., olive baboons, *Papio anubis*: Smuts and Watanabe, 1990; Guinea baboons, *Papio papio*: Whitham and Maestripieri, 2003; spotted hyenas, *Crocuta crocuta*: East et al., 1993; Tonkean macaques, *Macaca tonkeana*: De Marco et al., 2014). These affiliative interactions with intense physical contact require a high level of trust between involved individuals, similar to the ‘eye-poking’ ritual of Colombian white-faced capuchin (Cebus capucinus; Perry, 2011), and have been suggested to function as a signal to strengthen in-group affiliation commitment among party members, and to reinforcing social relationships. In birds, same-sex socio-sexual behaviours occur in over 130 species, and while the frequency of expression differs between species and often between the sexes of one species, it is related to parental care disparity between the sexes and the mating system; highly polygynous species with minimal male parental investment exhibit higher frequencies of male-male sexual behaviour (MacFarlane et al., 2010), whereas female-female sexual behaviour was reported more frequently in monogamous species (MacFarlane et al., 2010, 2007; Poiani, 2008). Non-conceptive male-female sexual behaviours in birds, for example copulations after clutch completion or outside of the fertile period, have been suggested to be important in the formation and maintenance of pair-bonds (social bond hypothesis; Birkhead et al., 1987). Less attention has been given to so-called reverse mountings in birds, *i.e*., a female mounting a male individual, although they have been described in approximately three dozen of species (reviewed in: Nuechterlein and Storer, 1989), including Northwestern crow (*Corvus caurin*; James, 1983), black-throated blue warbler (*Setophaga caerulescens; Marra, 1993)*, Rufous-tailed hawk *(Buteo ventralis; Raimilla et al., 2013)*, groove-billed ani (*Crotophaga sulcirostris*; Bowen et al., 1991), European shag (*Phalacrocorax aristotelis*; Ortega-Ruano and Graves, 1991), bearded vulture (*Gypaetus barbatus*; (Bertran and Margalida, 2006). Female-male mountings may be an integral part of courtship behaviour in monogamous, long-term bonded species, as described in different species of grebes (Nuechterlein & Storer, 1989). Interestingly, the frequency of female-male mountings may differ among pairs of the same species; in Dovekies (*Alle alle*) Jakubas and Wojczulanis-Jakubas (2008) noted that out of 851 copulations and copulation attempts observed among 37 pairs, only 5% were female-male mountings, whereas in one pair, half of the mountings were reversed. To what degree females play an active role in initiating these socio-sexual mountings and other behaviours that contribute to initiating or preventing divorce and maintain pair-bonds remains debated (Cézilly et al., 2000). More detailed data and meta-analytical approaches that consider heterogeneity among species, sexes, and individuals are needed to quantitatively assess the prevalence and systematically test hypotheses regarding the function of socio-sexual behaviours in birds.

Here, we report an observation of a female carrion crow mounting her pair-housed male partner. Carrion crows form long-term socially monogamous pairs (Glutz von Blotzheim, 1985; Meide, 1984), but the social system of carrion crows can be flexible, including territorial pairs, non-breeder flocks, and cooperatively breeding groups (Wascher, 2018). With the exemption of incubation and brooding, which is exclusively carried out by the female (Baglione and Canestrari, 2016), both sexes engage in parental care, *e.g*., nest building (Baglione & Canestrari, 2016), nest defence (Bossema and Benus, 1985), chick feeding (Canestrari et al., 2005), nest sanitation (Bolopo et al., 2015). Typical for non-tropical bird species, reproduction in carrion crows is seasonally limited to spring, when temperatures are mild and food availability high compared to other seasons (Ball and Ketterson, 2008). Reproduction is usually limited to one successful clutch a year, but replacement clutches may be laid after breeding failure (Roldán et al., 2013).

## Methods

### Study subjects

The female-male mounting we describe in detail below was recorded by CAFW, between two pair-housed crows in a population of captive carrion crows housed in a large outdoor aviary in Navafria, northern Spain (N 42°36’30 W 5°26’56; for further details on the population see: Wascher et al., 2019). At the time of the observation, crows were kept in four separate groups, two male-female pairs (hatched in 2010), one trio (two birds hatched in 2011, one bird hatched in 2007) and one group of seven juvenile individuals (hatched in 2012). Groups were visually but not acoustically separated from each other. The subjects of the reported observation hatched in 2010 and were taken from different nests in the wild between 11 and 22 days post hatching. Hence, the pair is considered unrelated. The subjects were hand-raised in captivity for the purpose of conducting studies on socio-cognitive abilities (Wascher et al., 2015) and social behaviour (Wascher et al., 2019). The crows were sexed using the P2/P8 molecular sexing method (Baglione and Canestrari, 2002), and separated into male-female pairs based on this information and behavioural observations (e.g., spatial proximity, low frequencies of aggressive behaviour, high frequencies of affiliative behaviour) in early 2012 (C. Núñez Cebrián, personal communication). As recommended by Webster and Rutz (2020) we explicitly declare and consider the STRANGEness of our study subjects: The observation has been reported in a captive pair of hand-raised individuals, i.e. the social background and rearing history differs from wild individuals. Social pairings have been decided by human observers based on social behaviour, which is different from mate choice in the wild. During the breeding season 2013, successful breeding occurred in another pair of the study population, who successfully raised two fledglings. The subjects of the present observation did not successfully breed in 2013 and 2014 and were not observed from 2015 onwards. Hand-raising and keeping crows in aviaries for non-invasive research on socio-cognitive behaviours were authorised by Junta de Castilla y Leon (núcleo zoologico 005074/2008; EP/LE/358/2009).

### Observation

Birds had been well habituated to the presence of CAFW since January 2013 and regular focal observations were conducted with caution to minimise disturbance to the focal individuals. Nesting material was provided to pair-housed birds during the breeding season, and nest checks were conducted from a distance, with a specialist mirror pole, when the female was off the nest.

The female-male mounting between two pair-housed crows was recorded during the breeding season in April 2013. The brief interaction was not filmed as part of systematic data collection but photographed by CAFW during animal care and aviary maintenance work using a Canon EOS 40D camera in burst mode. Using the continuous-shooting mode, a total of 24 pictures were taken at a rate of seven images per second, thus allowing for estimating of the duration of the mounting interaction to be approximately 3.5 seconds.

## Results

The observation reported here was made during the 2013 breeding season. On 26 ^th^ April 2013 in the early evening (approximately 18:30 CEST), CAFW observed the pair of carrion crows performing a courtship display, in which the female individual mounted the male individual (Figure 1). The photographs show the male individual crouching down, seemingly inviting the female to mount on top (Figure 1a-c). Based on the number of photographs taken during the mounting, the interaction was estimated to last approximately 3.5 seconds. Once mounted, the female performed a beak-touching display (Figure 1d-m). The pictures do not allow to clearly tell whether there is cloacal contact at any point of the interaction. After the mounting, both the male and the female engaged in a display similar to post-copulatory displays which CAFW had previously observed in common ravens (CAFW, personal observation), however the interaction was very short, and it is not well captured in the photographs (Figure 1o-q).

**Figure 1.**
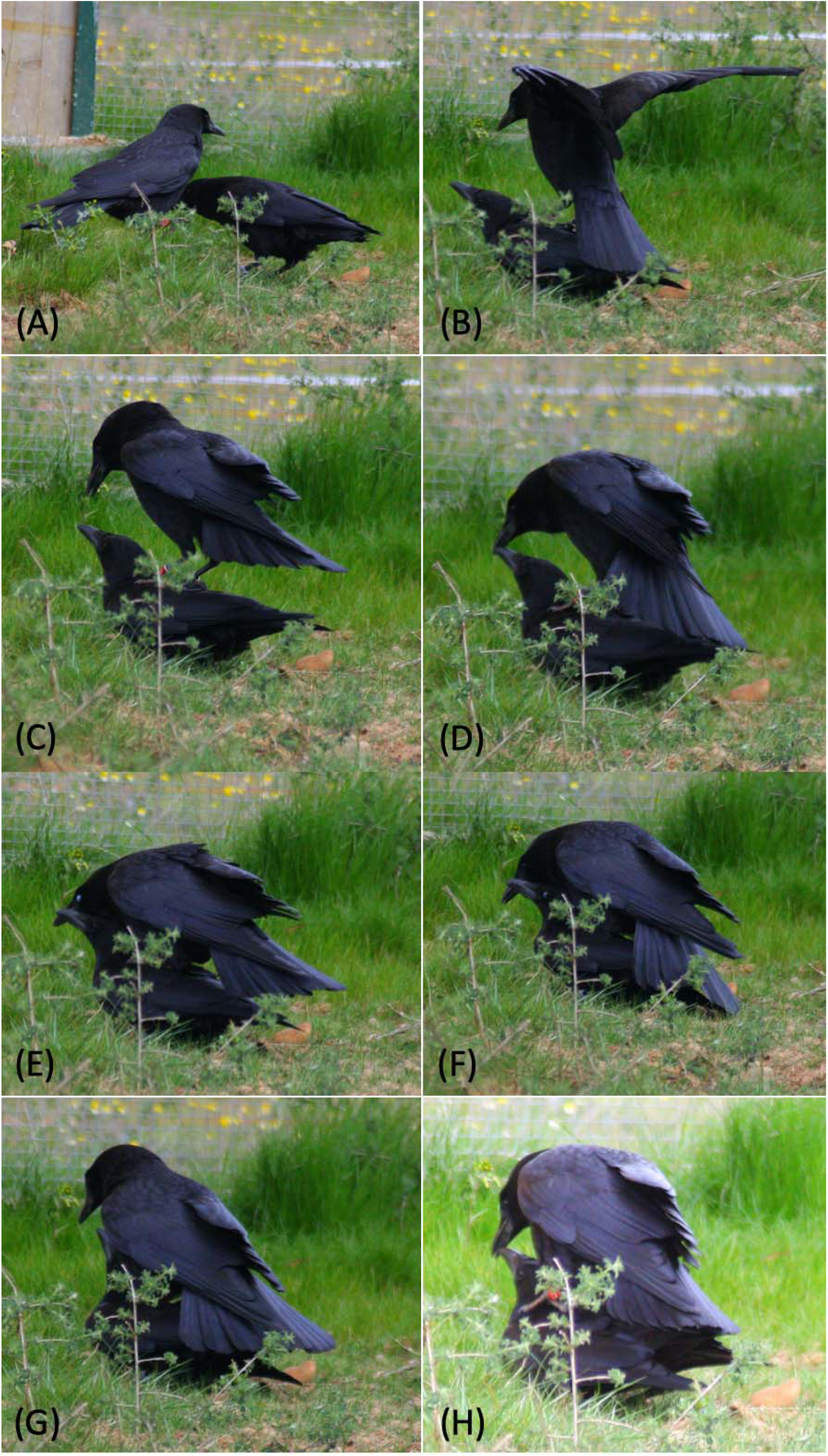
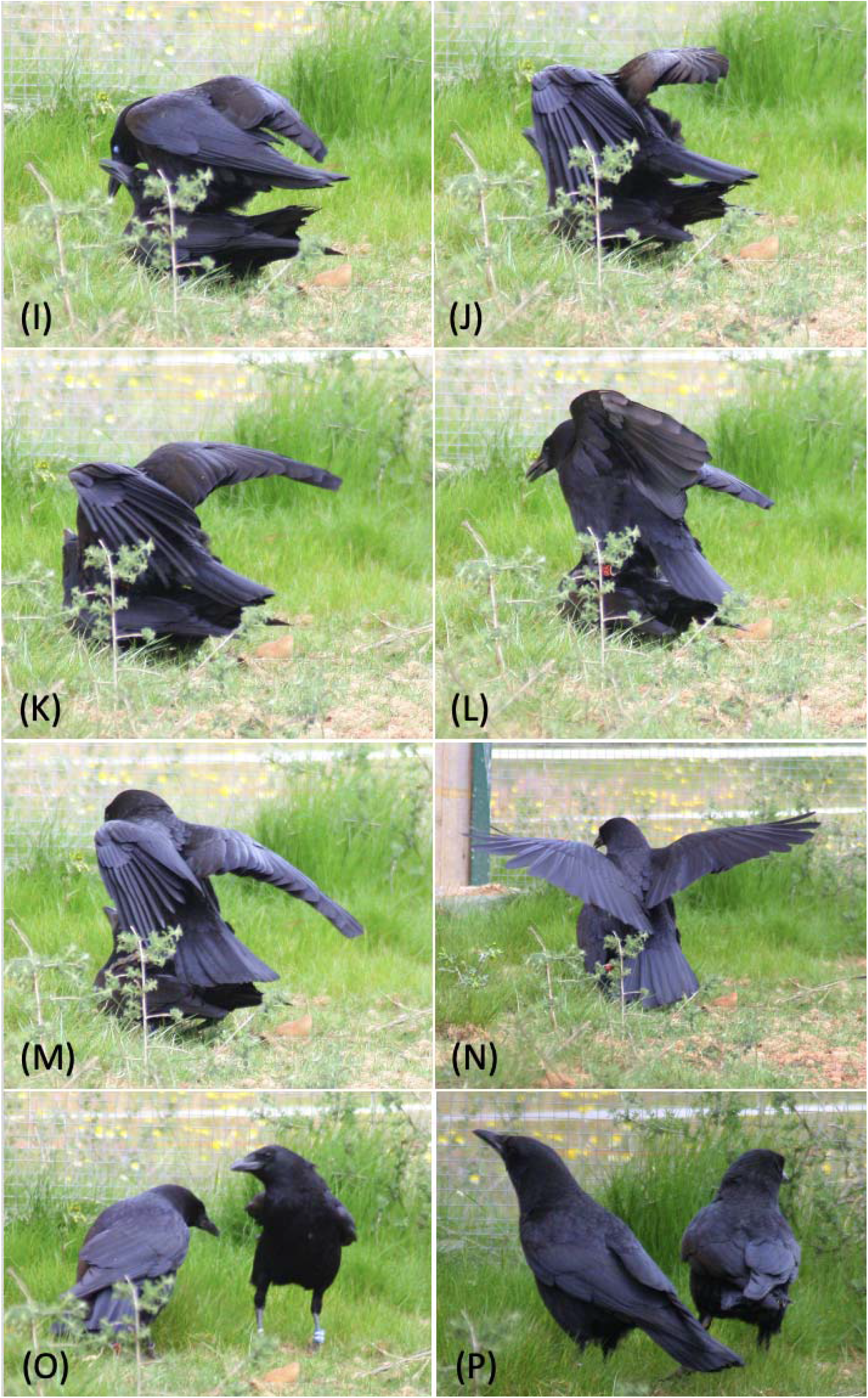
Series of 16 photographs taken by CAFW documenting an observation of a reversed, non-conceptive mounting on 26^th^ April 2013. A female carrion crow (red colour ring) is mounting a pair-bonded, male carrion crow (blue colour ring).

The focal individuals had two failed breeding attempts during the breeding season of 2013; eggs were found in their nest once prior and once after the observed reversed mounting. The female was repeatably reported on the nest since 15^th^ April 2013 and was assumed to have started incubation (CAFW and FH, personal observation). However, we did not confirm when the first egg was laid, as the female appeared unsettled and easily disturbed by the carers and other activity around the aviary, therefore no nest checks where conducted during that period. By 26^th^ April, when the reported observation was made, the female had abandoned the nest and eggshells were found on the ground of the aviary. The focal female was again regularly seen on the nest from 3^rd^ May onwards; the nest had two eggs on 6^th^ May and four eggs on 10^th^ May. On 14^th^ May, only one egg was left in the nest. The focal female was not observed on the nest anymore. A different pair of crows from the same captive population and of the same age as the focal pair successfully bred in 2013 and raised two chicks to fledging.

## Discussion

We report an observation of a female carrion crow mounting her co-housed male partner. Only few reports on frequency and variety of courtship behaviours (both non-conceptive mountings and male-female copulations) in corvids are published (Canestrari et al., 2022; Gill et al., 2020; Kilham, 1984; Townsend, 2009), including a previous report on female-male mounting in Northwestern crows (James, 1983). Sexual behaviours and copulations are brief among pairs of crows and other corvids, may occur both on the ground and in trees, and -- unless individually marked and sexed -- in these monomorphic species it is difficult to determine the sex of the involved individuals. We estimated the female-male mounting to last for approximately three to four seconds. A previous review of data and grey literature concerning avian copultation behaviour in 131 species found that copulations lasted less than five seconds in a third of species for which duration information was avialable (69 species), and less than ten seconds in over 55% of those species (Birkhead et al., 1987). Empirical data on reproductive and socio-sexual behaviours are therefore difficult to gather and virtually lacking. Only few studies with high observational effort report copulatory behaviours among corvids, for example in American crows (Kilham, 1984; Townsend, 2009)and carrion crows (Canestrari et al., 2022), and, Gill et al. (2000) used full-day nest-box video surveillance and microphone backpacks to directly observe sexual behaviour of jackdaws (*Corvus monedula (Gill et al., 2020)*) and In our captive study population, in approximately 100 hours of focal recordings between 2013 and 2014 and many more hours of behavioural observations during experiments and animal keeping duties or habituation of crows, no other incidents of courtship behaviours or mating had been observed.

Data on both copulations and resulting parentships, and non-conceptive socio-sexual behaviours are important to test hypotheses about potential evolutionary drivers of non-conceptive sexual behaviour (Bailey and Zuk, 2009). For example, female-male mountings might be a regular element of crow courtship behaviour, similar to displays such as bowing, billing, and vocal exchanges that are part of courtship and pair formation behaviours (*e.g*., American crows: Kilham 1985) or these non-conceptive mountings may play a particular role in maintaining and strengthening the pair-bond (Birkhead et al., 1987), for example after a failed breeding attempt as reported here.

The focal pair has never successfully raised offspring together, neither in the year of the observation, or before or afterwards. Captive conditions such as disturbance by other crows in neighbouring aviary compartments may have contributed to unsuccessful reproductive attempts however, we note that other pairs of crows and other corvid species have repeatedly successfully bred in captivity, including a pair of crows from the same population and cohort, that successfully raised chicks in the breeding season of 2013. Hand-raising corvids for captive populations is a common practice in comparative cognitive research. While it might have effects on the development of the individuals’ socio-sexual behaviour, we are not aware of any other reports of female-male mounting or other socio-sexual behaviours perceived as unusual in captive corvids. Our anecdotal report therefore remains a valid observation and we hope to stimulate more systematic investigations of this behaviour both in captive populations and in the wild. We highlight the importance of more systematic investigations of rare social-sexual behaviours, which might potentially hold insights into an understanding of evolutionary drivers of non-conceptive sexual behaviours.

## Acknowledgements

We are grateful to Prof. Vittorio Baglione and Prof. Daniela Canestrari for access to the study population and general support and discussion.

## References

Baglione, V., Canestrari, D., 2016. Carrion crows: Family living and helping in a flexible social system., in: Cooperative Breeding in Vertebrates: Studies of Ecology, Evolution, and Behavior. Cambridge University Press, Cambridge, pp. 97–114.

Baglione, V., Canestrari, D., 2002. Cooperatively breeding groups of carrion crow (Corvus corone corone) in northern Spain. The Auk 119, 790–799. 10.2307/4089974

Bailey, N.W., French, N., 2012. Same-sex sexual behaviour and mistaken identity in male field crickets, Teleogryllus oceanicus. Anim. Behav. 84, 1031–1038. 10.1016/j.anbehav.2012.08.001

Bailey, N.W., Zuk, M., 2009. Same-sex sexual behavior and evolution. Trends Ecol. Evol. 24, 439–446. 10.1016/j.tree.2009.03.014

Ball, G.F., Ketterson, E.D., 2008. Sex differences in the response to environmental cues regulating seasonal reproduction in birds. Philos. Trans. R. Soc. B Biol. Sci. 363, 231–246. 10.1098/rstb.2007.2137

Bertran, J., Margalida, A., 2006. Reverse mounting and copulation behavior in polyandrous bearded vulture (Gypaetus barbatus) trios. Wilson J. Ornithol. 118, 254–256. 10.1676/05-046.1

Birkhead, T.R., Atkin, L., Møller, A.P., 1987. Copulation behaviour of birds. Behaviour 101, 101–138. 10.1163/156853987X00396

Bolopo, D., Canestrari, D., Marcos, J.M., Baglione, V., 2015. Nest sanitation in cooperatively breeding Carrion crows. The Auk 132, 604–612. 10.1642/AUK-14-233.1

Bossema, I., Benus, R.F., 1985. Territorial defence and intra-pair cooperation in the carrion crow (Corvus corone). Behav. Ecol. Sociobiol. 16, 99–104. 10.1007/BF00295141

Bowen, B.S., Koford, R.R., Vehrencamp, S.L., 1991. Seasonal pattern of reverse mounting in the groove-billed ani (Crotophaga sulcirostris). The Condor 93, 159–163. 10.2307/1368618

Canestrari, D., Marcos, J., Baglione, V., 2005. Effect of parentage and relatedness on the individual contribution to cooperative chick care in carrion crows Corvus corone corone. Behav. Ecol. Sociobiol. 57, 422–428. 10.1007/s00265-004-0879-1

Canestrari, D., Trapote, E., Vila, M., Baglione, V., 2022. Copulations with a nestling by an adult care-giver in a kin-living bird. Behaviour 160, 191–199. 10.1163/1568539X-bja10195

Cézilly, F., Préault, M., Dubois, F., Faivre, B., Patris, B., 2000. Pair-bonding in birds and the active role of females: a critical review of the empirical evidence. Behav. Processes 51, 83–92. 10.1016/S0376-6357(00)00120-0

Dal Pesco, F., Fischer, J., 2018. Greetings in male Guinea baboons and the function of rituals in complex social groups. J. Hum. Evol. 125, 87–98. 10.1016/j.jhevol.2018.10.007

De Marco, A., Sanna, A., Cozzolino, R., Thierry, B., 2014. The function of greetings in male Tonkean macaques. Am. J. Primatol. 76, 989–998. 10.1002/ajp.22288

East, M.L., Hofer, H., Wickler, W., 1993. The erect “penis” is a flag of submission in a female-dominated society: greetings in Serengeti spotted hyenas. Behav. Ecol. Sociobiol. 33. 10.1007/BF00170251

Faraut, L., Northwood, A., Majolo, B., 2015. The functions of non-reproductive mounts among male Barbary macaques (Macaca sylvanus): Socio-Sexual Mounts in Barbary Macaques. Am. J. Primatol. 77, 1149–1157. 10.1002/ajp.22451

Gill, L.F., Van Schaik, J., Von Bayern, A.M.P., Gahr, M.L., 2020. Genetic monogamy despite frequent extrapair copulations in “strictly monogamous” wild jackdaws. Behav. Ecol. 31, 247–260. 10.1093/beheco/arz185

Glutz von Blotzheim, U.N., 1985. Handbuch der Vögel Mitteleuropas. Aula-Verlag, Wiesbaden.

Gómez, J.M., Gónzalez-Megías, A., Verdú, M., 2023. The evolution of same-sex sexual behaviour in mammals. Nat. Commun. 14, 5719. 10.1038/s41467-023-41290-x

Gunst, N., Casarrubea, M., Vasey, P.L., Leca, J.-B., 2020. Is female-male mounting functional? An analysis of the temporal patterns of sexual behaviors in Japanese macaques. Physiol. Behav. 223, 112983. 10.1016/j.physbeh.2020.112983

Ham, J.R., Malin, L.K., Manitzas Hill, H.M., 2023. Non-conceptive sexual behavior in cetaceans: comparison of form and function., in: Sex in Cetaceans. Springer, Texas.

Jakubas, D., Wojczulanis-Jakubas, K., 2008. Variation in copulatory behavior of the Dovekie. Waterbirds 31, 661–665. 10.1675/1524-4695-31.4.661

James, P.C., 1983. Reverse mounting in the Northwestern Crow. J. Field Ornithol. 54, 418.419.

Kilham, L., 1984. Cooperative breeding of American crows. J. Field Ornithol. 55, 349–356.

Kotrschal, K., Hemetsberger, J., Weiss, B.M., 2006. Making the best of a bad situation: homosociality in greylag geese., in: Homosexual Behaviour in Animals: An Evolutionary Perspective. Cambridge University Press, Cambridge, pp. 45–76.

Lilley, M.K., Ham, J.R., Hill, H.M., 2020. The development of socio-sexual behavior in belugas (Delphinapterus leucas) under human care. Behav. Processes 171, 104025. 10.1016/j.beproc.2019.104025

Macchiano, A., Razik, I., Sagot, M., 2018. Same-sex courtship behaviors in male-biased populations: evidence for the mistaken identity hypothesis. Acta Ethologica 21, 147–151. 10.1007/s10211-018-0293-8

MacFarlane, G.R., Blomberg, S.P., Kaplan, G., Rogers, L.J., 2007. Same-sex sexual behavior in birds: expression is related to social mating system and state of development at hatching. Behav. Ecol. 18, 21–33. 10.1093/beheco/arl065

MacFarlane, G.R., Blomberg, S.P., Vasey, P.L., 2010. Homosexual behaviour in birds: frequency of expression is related to parental care disparity between the sexes. Anim. Behav. 80, 375–390. 10.1016/j.anbehav.2010.05.009

Manson, J.H., Perry, S., Parish, A.R., 1997. Nonconceptive sexual behavior in bonobos and capuchins. Int. J. Primatol. 18, 767–786.

Marra, P.P., 1993. Reverse mounting in the Black-throated Blue Warbler. Wilson Bull. 105, 359–361.

Meide, M., 1984. Raben- und Nebelkrähe. Westarp Wissenschaften, Magdeburg.

Nuechterlein, G.L., Storer, R.W., 1989. Reverse mounting in grebes. The Condor 91, 341. 10.2307/1368312

Ortega-Ruano, J., Graves, J.A., 1991. Reverse mounting during the courtship of the European shag Phalacrocorax aristotelis. The Condor 93, 859. 10.2307/3247720

Perry, S., 2011. Social traditions and social learning in capuchin monkeys (Cebus). Philos. Trans. R. Soc. B Biol. Sci. 366, 988–996. 10.1098/rstb.2010.0317

Poiani, A., 2008. Same-sex mounting in birds: Comparative test of a synthetic reproductive skew model of homosexuality. Open Ornithol. J. 1, 36–45. 10.2174/1874453200801010036

Raimilla, V., Norambuena, H.V., Jiménez, J.E., 2013. A record of reverse mounting in the Rufoustailed hawk (Buteo ventralis) in Southern Chile. J. Raptor Res. 47, 326–327. 10.3356/JRR-12-47.1

Roldán, M., Martín-Gálvez, D., Rodríguez, J., Soler, M., 2013. Breeding biology and fledgling survival in a carrion crow (Corvus corone) population of southern spain: A comparison of group and pair breeder. Acta Ornithol. 48, 221–235. 10.3161/000164513X678865

Smuts, B.B., Watanabe, J.M., 1990. Social relationships and ritualized greetings in adult male baboons (Papio cynocephalus anubis). Int. J. Primatol. 11, 147–172. 10.1007/BF02192786

Townsend, A.K., 2009. Extrapair copulations predict extrapair fertilizations in the American crow. The Condor 111, 387–392. 10.1525/cond.2009.090010

Wascher, C.A.F., 2018. Corvids, in: Vonk, J., Shackelford, T. (Eds.), Encyclopedia of Animal Cognition and Behavior. Springer International Publishing, Cham, pp. 1–12. 10.1007/978-3-319-47829-6_1799-1

Wascher, C.A.F., Canestrari, D., Baglione, V., 2019. Affiliative social relationships and coccidian oocyst excretion in a cooperatively breeding bird species. Anim. Behav. 158, 121–130. 10.1016/j.anbehav.2019.10.009

Wascher, C.A.F., Hillemann, F., Canestrari, D., Baglione, V., 2015. Carrion crows learn to discriminate between calls of reliable and unreliable conspecifics. Anim. Cogn. 18, 1181–1185. 10.1007/s10071-015-0879-8

Webster, M.M., Rutz, C., 2020. How STRANGE are your study animals? Nature 582, 337–340. 10.1038/d41586-020-01751-5

Whitham, J.C., Maestripieri, D., 2003. Primate rituals: The function of greetings between male Guinea baboons. Ethology 109, 847–859. 10.1046/j.0179-1613.2003.00922.x

